# Insect-specific Yada Yada virus chimeric vaccines protect against chikungunya and Ross River virus-induced arthritis

**DOI:** 10.1101/2025.07.28.667096

**Authors:** Wilson Nguyen, Agnes Carolin, Mikaela G. Bell, Bing Tang, Kexin Yan, Abigail L. Cox, Andreas Suhrbier, Jessica J. Harrison, Jody Hobson-Peters, Daniel J. Rawle

## Abstract

Arthritogenic alphaviruses such as chikungunya virus (CHIKV) and Ross River virus (RRV) are mosquito-borne viruses that can cause debilitating polyarthritis/polyarthralgia in humans. Although two CHIKV vaccines have been licensed, there are no licensed vaccines for RRV. Herein we generate a host-restricted, insect-specific alphavirus, Yada Yada virus (YYV), chimeric vaccine for CHIKV (YYV-CHIKV_Mauritius_) and for RRV (YYV-RRV_TT_). YYV-CHIKV_Mauritius_ and YYV-RRV_TT_ was able to replicate in C6/36 mosquito cells to similar titres as wild-type CHIKV and RRV. YYV-CHIKV_Mauritius_ was also neutralised by CHIKV monoclonal antibodies to the same titres as wild-type CHIKV, indicating its potential as a diagnostic antigen to detect neutralising CHIKV antibodies in human or animal sera. YYV-CHIKV_Mauritius_ further demonstrated protection against CHIKV infection and disease in a wild-type mouse model. Two doses of YYV-CHIKV_Mauritius_ showed anti-CHIKV ELISA and neutralising antibody responses, with protection against foot swelling, viraemia and viral feet tissue titres. Protection against CHIKV histopathology including myositis, tendonitis, arthritis, subcutaneous oedema and haemorrhage was also observed. YYV-RRV_TT_ also demonstrated protection against RRV infection and disease in a wild-type mouse model, with two vaccine doses inducing anti-RRV ELISA and neutralising antibody responses. Protection against foot swelling, viraemia and viral feet tissue titres and RRV histopathology including myositis, tendonitis, arthritis and subcutaneous oedema was also observed. Cross-protection was also evaluated between YYV-CHIKV_Mauritius_ and RRV. Although cross-reactive total IgG were observed for YYV-CHIKV_Mauritius_ vaccinated mice, this offered no cross-neutralising antibodies and no protection against RRV infection and disease. Overall, our findings show that YYV-CHIKV_Mauritius_ and YYV-RRV_TT_ are safe and efficacious vaccines against CHIKV and RRV, respectively, but do not offer cross-protection.

## 1. Introduction

Alphaviruses are small, enveloped, positive-sense single-stranded RNA viruses in the *Togaviridae* family, with the alphavirus genus comprising over 30 distinct members that are classified into seven antigenic serogroup complexes. Primarily transmitted by mosquitoes, alphaviruses cause rheumatic disease in humans and are globally distributed. Arthritogenic alphaviruses such as chikungunya virus (CHIKV), Ross River virus (RRV), Mayaro virus (MAYV), Sindbis (SINV) and Barmah Forest virus (BFV) cause a variety of disease manifestations in humans, with severe cases experiencing polyarthritis/polyarthralgia (Nguyen et al., 2020).

CHIKV continues to cause sporadic outbreaks, with the largest recorded epidemic beginning in 2004 and spreading to more than 100 countries over four continents, resulting in >10 million cases of debilitating rheumatic disease and mortality estimates ranging from 0.024%-0.7% (Suhrbier, 2019). In comparison, RRV causes ≈4600 cases annually in Australia, where it is a notifiable disease (Farmer and Suhrbier, 2019). RRV is endemic to Australia and parts of Papua New Guinea, with previous largescale outbreaks occurring in the Pacific Islands in 1979/1980s (Harley et al., 2001). Serological studies have also indicated silent circulation of RRV in regions of French Polynesia and American Samoa (Lau et al., 2017). Recent outbreaks in the Australian Shoalwater Bay Military Training Area, also highlights the potential risk of international dissemination (Shanks, 2019). RRV is estimated to cost ∼$15 million per year in terms of health care and lost productivity (Harley et al., 2001; Koolhof et al., 2020), with a burden of 35.8 years lived with disability (YLDs) between 2003-2018 (Damtew et al., 2024). While a formalin and UV-inactivated whole virus, alum-adjuvanted RRV vaccine passed phase III clinical trials and was well tolerated and immunogenic (Wressnigg et al., 2015), the vaccine was never commercialised, most likely due to the relatively low recognised case numbers and high costs associated with bringing a vaccine to the market.

Despite the continued worldwide burden of CHIKV, it was only recently that a licensed vaccine (IXCHIQ®) was made available for human use (U.S. Food & Drug Administration, 2023). IXCHIQ® is a live-attenuated, single-dose vaccine approved for use in adults 18 years or older and is recommended for adults travelling to areas with CHIKV outbreaks (U.S. Centers for Disease Control and Prevention, 2025). In May of 2025, a second CHIKV vaccine from Bavarian Nordic A/S (VIMKUNYA™) was FDA approved (Bavarian Nordic, 2025). VIMKUNYA™ is a virus-like particle single-dose vaccine recommended for use in people 12 years or older, and to those travelling to areas prone to CHIKV outbreaks (U.S. Centers for Disease Control and Prevention, 2025).

Prior to the licensing of IXCHIQ® and VIMKUNYA™, several CHIKV candidates were evaluated at various pre-clinical and clinical trials. Nakao & Hotta (1973) showed that a UV-inactivated whole virus CHIKV was immunogenic in monkeys. A formalin-inactivated CHIKV vaccine demonstrated immunogenicity in mice stimulating humoral and cell-mediated immune responses (Tiwari et al., 2009). A DNA vaccine expressing the CHIKV capsid and envelope protein sequences showed high antibody responses in C57BL/6J mice (Muthumani et al., 2008). Protein sub-unit vaccines expressing envelope proteins (Kumar et al., 2012) and virus-like particle vaccines (VLPs) (Metz et al., 2013) showed protection against CHIKV challenge in mice models. A live attenuated CHIKV vaccine, TSI-GSD-218 (U.S. Army Medical Research Institute for Infectious Diseases), was evaluated in phase II human trials, but was associated with arthralgia (Edelman et al., 2000). Bharat Biotech (Hyderabad, India) showed immunogenicity in monkeys with a virus-like particle (Cherian et al., 2023). An inactivated, alum adjuvanted vaccine, BBV87 (International Vaccine Institute), and a VLP vaccine, PXVX0317 (Emergent BioSolutions), are currently in late-phase development (phase II/III). Chimeric vaccines for CHIKV have also been developed using the Venezuelan equine encephalitis virus (VEEV) TC-83 vaccine strain, or Sindbis virus as a backbone with the structure protein genes of CHIKV substituted. These chimeric vaccines demonstrated robust neutralising antibodies responses with complete protection against disease and viraemia in mice (Wang et al., 2008).

Chimeric vaccines offer unique advantages over other types of vaccines due to their hybrid design, scalability, and immunogenicity, mimicking the antigenic structure of wild-type viruses. Insect-specific viruses (ISV) chimeric vaccines offer an additional advantage in which they can only replicate in insect cells and cannot produce viral progeny in mammalian cells, yet can stimulate immune responses in vertebrate hosts. Eilat (EILV) is an insect-specific alphavirus (ISA) which has been exploited to generate an EILV/CHIKV vaccine containing the non-structural proteins (nsP1, nsP2, nsP3 and nsP4) of EILV and the structural proteins of CHIKV (E1, 6K, E2, E3 and capsid). A single unadjuvanted dose of 10^8.1^ PFU of EILV/CHIKV-sP (live virus, PEG-precipitated) protected mice and non-human primates from CHIKV challenge (Adam et al., 2024).

Herein, we used a recombinant vaccine platform based on the insect-specific alphavirus Yada Yada virus (YYV) which we have previously demonstrated to have a robust safety profile and mimics the virion antigenic structure for a range of pathogenic alphaviruses (Bell et al., 2024). We constructed a chimeric vaccine that authentically displays the immunogenic structural proteins (Capsid, E3, E2, 6K and E1) of CHIKV_Mauritius_ and RRV_TT_. These chimeric vaccines were shown to be effective and safe vaccines against CHIKV_Reunion_ and RRV_TT_ challenge, respectively, in mouse models of infection and disease. We also evaluated cross-protection of CHIKV_Mauritius_ against RRV challenge in mice.

## 2. Methods

### 2.1. Ethics Statement and Approvals

All mouse work was conducted in accordance with the “Australian code for the care and use of animals for scientific purposes” as defined by the National Health and Medical Research Council of Australia. Mouse work performed at QIMR Berghofer was approved by the QIMR Berghofer Animal Ethics Committee (P3746, A2108-612). All CHIKV work was conducted in a biosafety level 3 (BSL3) facility at the QIMR Berghofer (Australian Department of Agriculture, Fisheries and Forestry certification Q2326 and Office of Gene Technology Regulator certification 3445). All work was approved by the QIMR Berghofer Safety Committee (P3746). Use of genetically modified chimeric viruses at QIMR Berghofer was approved by the Gene Technology Sub-Committee with the identifier GTSC_077_2025: NLRD 2.1(d). All work with infectious and circular polymerase extension reaction (CPER)-generated viruses was approved by the UQ Institutional Biosafety Committee (UQ IBC, approvals IBC/1246/SCMB/2019, IBC/1389/SCMB/2021, and IBC/1402/SCMB/2022).

### 2.2. Cell lines and culture

C6/36 (*Aedes albopictus*) cells (ATCC# CRL-1660) were maintained in Roswell Park Memorial Institute 1640 (RPMI 1640) medium (Thermo Fisher Scientific, Scoresby, VIC, Australia), supplemented with 5-10% foetal bovine serum (FBS) (Sigma-Aldrich, Castle Hill, NSW, Australia) at 27-28 °C. Vero (*Cercopithecus aethiops*, African green monkey kidney) cells (ATCC# CCL-81) were maintained in RPMI 1640, supplemented with 10% FBS at 37 °C with 5% CO_2_. All media were supplemented with 50 U/ml penicillin, 50 µg/mL streptomycin and 2 mmol/L L-glutamine (PSG).

### 2.3. Virus isolates and culture

The Réunion Island CHIKV isolate (LR2006-OPY1) is a primary human isolate from the 2012 outbreak in Réunion Island and was passaged twice in C6/36 cells (GenBank: KT449801). The Mauritius CHIKV isolate (GenBank: MH229986) is a primary human isolate from the 2006 outbreak in Mauritius passaged once in C6/36 cells. RRV_TT_ (GenBank: KY302801.2) is a primary human isolate from the 2014 Australian transfusion transmission case (Hoad et al., 2015) passaged twice in C6/36 cells. Virus stocks were generated by infecting C6/36 monolayers with virus as described in (Bell et al., 2024; Gardner et al., 2010; Nguyen et al., 2020). Virus titres were determined by an endpoint dilution where cytopathic effects were evident, and titres (CCID_50_) were calculated based on the method of Spearman and Karber (Ramakrishnan, 2016) (a conventional Excel CCID calculator is available at https://www.klinikum.uni-heidelberg.de/zentrum-fuer-infektiologie/molecular-virology/welcome/downloads).

### 2.4. Infectious clone and YYV-CHIKV_Mauritius_ chimeric virus generation using circular polymerase extension reaction (CPER)

An infectious DNA construct was created using the circular polymerase extension reaction (CPER) method as described (Bell et al., 2024). In brief, cDNA was synthesised from extracted viral RNA using Superscript IV Reverse Transcriptase (Invitrogen) using an Oligo(dt) primer, random hexamers or a virus-specific reverse primer as per manufacturer’s instructions. cDNA was used as a template to generate overlapping dsRNA PCR fragments containing 15-30 nucleotide overhangs at both 5’ and 3’ ends using Q5 High-Fidelity DNA polymerase (New England Biolabs) as per manufacturer’s instructions. The structural protein cassettes as dsDNA were generated for CHIKV_Mauritius_ using GenBank MH229986 sequence data and the following primers: Capsid (Forward: 5’-CCAACCAGCCTAACCATGGAGTTCATCCCAACCC-3’; Reverse: 5’-GGATGAACTCCATGGTTAGGCTGGTTGGGGTAG-3’) and envelope protein 1 (E1) (Forward: 5’-CGTGTCGTTCAGCAGGCACTAATCCACTTATAGCACTATAG-3’; Reverse: 5’-CTATAGTGCTATAAGTGGATTAGTGCCTGCTGAACGACACG-3’).

### 2.5. YYV-RRV_TT_ and YYV-CHIKV_Mauritius_ vaccine production

The YYV-CHIKV_Mauritius_ and YYV-RRV_TT_ vaccines were generated as previously described (Bell et al., 2024). In brief, Equimolar amounts (0.1 pmol) of each viral cDNA fragment, together with a linker region (amplified with virus-specific overhangs from plasmid DNA) containing a modified insect promoter and hepatitis delta virus ribozyme as previously described (Amarilla et al., 2021), were added to a Q5 PCR reaction as per the manufacturer’s instructions and cycled under the following conditions: 98 °C for 2 min (1 cycle), 98 °C for 30 s, 55 °C for 30 s, 72 °C for 6 min (2 cycles), 98 °C for 30 s, 55 °C for 30 s, 72 °C for 8 min (10 cycles). The entire reaction was transfected using TransIT-LT1 Transfection Reagent (Mirus) as per the manufacturer’s instructions and incubated for 5–7 dpi. Cells and virus supernatant (P0) were passaged onto freshly seeded C6/36 cell monolayers for IFA analysis as described below to confirm the presence of successful CPER virus recovery. Coverslips were stained with virus-specific E1/E2 mAbs, or anti-dsRNA mAbs (3G1/2G4) if no mAbs were available for the target virus. Virus identity was confirmed by RT-PCR of RNA extracted from cell culture supernatant. Sequences of chimeras was confirmed via Sanger sequencing at the Australian Genome Research Facility (AGRF, Brisbane, Australia).

Both vaccines were purified to the sucrose cushion level (YYV-RRV_TT_) or potassium tartrate level (YYV-CHIKV_Mauritius_) as previously described (any of our references). Briefly, sub-confluent monolayers of C6/36 cells were infected with chimeric virus at an MOI of 0.1. The supernatant was collected and 3-, 5- and 7-dpi. The virus culture supernatant was clarified by centrifugation at 3000 rpm for 30 min at 4 °C, before filtering through a 0.22 µM filter. After each collection, cells were replenished with fresh RPMI containing 2% FBS. The virions were precipitated via the addition of polyethylene glycol 8000 to a final concentration of 8% and slow stirring overnight at 4 °C. The virus was pelted at 8000 rpm (Beckman Coulter JLA 10.500 rotor) for 1.5 h at 4 °C, before ultracentrifugation through a sucrose cushion and a potassium tartrate gradient as described previously. The YYV-RRV_TT_ chimera was not taken through the potassium tartrate gradient, as sucrose cushion resulted in sufficient sample purity, due to minimal production of CPE. The purified virus was collected, and buffer exchanged into sterile PBS using a 30 kDa molecular weight cut-off amicon filter and stored at 4 °C. Purified virion proteins were resolved by SDS-PAGE (NuPAGE 4 -12% Bis-Tris gels, Invitrogen) and staining with SYPRO Ruby stain (Invitrogen) as per the manufacturer’s instructions.

### 2.6. Mice, infection and disease evaluation vaccination and CHIKV challenge

Mice were housed under the following conditions as described (Abbo et al., 2023; Nguyen et al., 2020; Rawle et al., 2020): 12:12 light-dark cycle; 7:45 a.m. sunrise and 7:45 pm sunset; 15-min light-dark dark-light ramping times. Enclosures were M.I.C.E cages (Animal Care Systems, Centennial, CO, USA). Ventilation was with 100% fresh air, 8 complete air exchanges/h/room. Temperature was 22 ± 1°C. In-house enrichment used paper cups, tissue paper, and cardboard rolls. Bedding was PuraChips (Able Scientific, Perth, WA, Australia) (aspen fine). Food was double-bagged Norco rat and mouse pellet (AIRR, Darra, QLD, Australia). Water was deionized water acidified with HCl (pH = 3.2).

Mouse models for CHIKV have been established as described (Gardner et al., 2010; Metz et al., 2013; Nguyen et al., 2020; Poo et al., 2014) using the CHIKV_Reunion_ isolate. CHIKV_Reunion_ and CHIKV_Mauritius_ belong in the same phylogenetic clade with 100% amino acid identity for the structural proteins between the two isolates. Thus, the CHIKV_Reunion_ isolate was chosen as the challenge isolate for this study. Mouse models for RRV have been established as described (Nguyen et al., 2020; Rawle et al., 2020) using the human isolate RRV_TT_.

In brief, mice were infected with 4 log_10_ CCID_50_ of CHIKV_Reunion_ or RRV_TT_ subcutaneously into the top or side of each hind foot as described (Gardner et al., 2010; Nguyen et al., 2020). Serum viraemia, foot swelling, tissue titres and histology were evaluated as previously described (Gardner et al., 2010; Nguyen et al., 2020; Poo et al., 2014; Wilson et al., 2017).

### 2.7. Mice vaccination and challenge

Female C57BL/6J (6 week old, 12 per group) mice were purchased from OzGene (Canning Vale, Western Australia, Australia). Mice were vaccinated twice with either 1 µg purified YYV-CHIKV_Mauritius_ or YYV-RRV_TT_, or PBS (without adjuvant) separated by ∼4 weeks, as described previously (Abbo et al., 2023; Miao et al., 2024; Nguyen et al., 2020; Rawle et al., 2020). Briefly, vaccines were administered intramuscularly (i.m.) into both quadriceps muscles of anaesthetised mice with 50 µl per muscle using a 27 G needle. Serum neutralising antibodies and ELISA responses were determined after the first vaccination (∼4 weeks), and booster vaccination (∼8 weeks). Six weeks after the second vaccination, mice were challenged with 10^4^ CCID_50_ diluted to a volume of 40 µl of CHIKV_Reunion_ or RRV_TT_ subcutaneously into the top or side of each hind foot as described (Gardner et al., 2010; Nguyen et al., 2020).

### 2.8. End-point antibody ELISA and neutralisation titre determination

Blood was collected at the indicated time points via lateral tail vein bleed into serum separator tubes and the aliquoted serum was stored at – 20°C. IgG responses were determined by standard ELISA using whole CHIKV_Reunion_ or RRV_TT_ as antigen. The antigen was purified from infected C6/36 cell supernatants by 40% PEG-6000 precipitation (Sigma-Aldrich, St. Louis, MO, USA) and ultracentrifugation (Beckman floor standing ultra, Beckman Coulter, CA, USA) at ∼134,000 rcf at 4°C for 2 h through a 20% sucrose (Sigma-Aldrich, St. Louis, MO, USA) cushion. Endpoint ELISA titres were determined as described previously (Gardner et al., 2010; Nguyen et al., 2020). Briefly, serum samples, starting at a 1:30 dilution, were serially diluted 1:2 in duplicate and bound antibody detected using biotin-labelled rat anti-mouse-IgG (Thermo Fisher Scientific), streptavidin HRP (Biosource, Camarillo, CA, USA), and 2,2′-azinobis(3-ethylbenzothiazolinesulfonic acid) substrate (Sigma-Aldrich, St. Louis, MO, USA). Endpoint titres were interpolated when optical density at 405 nm (OD_405_) values reached the mean OD_405_ + 3 standard deviations for naive serum.

Neutralisation assays were performed as previously described (Abbo et al., 2023; Nguyen et al., 2020; Rawle et al., 2020). Briefly, mouse serum samples were heat-inactivated (56°C for 30 min) and incubated in duplicate with 200 CCID_50_ of CHIKV_Reunion_ or RRV_TT_ at 37°C for 1 h before Vero cells were added at a concentration of 10^5^ cells/well. The initial serum dilution was 1:40 or 1:50 for RRV and CHIKV, respectively, with serial dilutions of 1:2 in duplicate. After 5-7 days, cells were fixed and stained with formaldehyde and crystal violet, and the 50% neutralising titres were interpolated from optical density (OD_590_) values versus serum dilution plots, as described (Abbo et al., 2023; Nguyen et al., 2020; Rawle et al., 2020).

### 2.9. Alphavirus challenge and disease determination

Blood was collected at the indicated time points via lateral tail vein bleed into serum separator tubes and the aliquoted serum was stored at – 20°C. Six mice per group were euthanised on day 6 or day 7 post-challenge (peak foot swelling time point) and feet were harvested and collected. Tissue and serum titres were determined by CCID_50_ assays as previously described (Gardner et al., 2010; Nguyen et al., 2020). For serum titrations, samples were titrated in duplicate starting with a 1 in 10 dilution followed by a 2-fold serial dilution on C6/36 cells (3-day culture). Subsequently, parallel well-to-well transfer of supernatants into 96-well plates containing Vero cells was carried out. After 3 days of culture, cytopathic effects were observed. For tissue titrations, samples were titrated in quadruplicate starting with a 1 in 10 dilution followed by a 2-fold serial dilution on C6/36 cells (3-day culture), followed by a parallel transfer onto Vero cells, with cytopathic effects observed after 3 days of culture.

Foot swelling (height and width of the perimetatarsal area) of the hind feet were measured using Kincrome digital vernier calipers as previously described (Abbo et al., 2023; Gardner et al., 2010; Nguyen et al., 2020; Poo et al., 2014; Rawle et al., 2020).

### 2.10. Histology

Histology and quantitation of staining were undertaken as described previously (Gardner et al., 2010; Nguyen et al., 2020; Rawle et al., 2020). In brief, feet were fixed in 10% formalin, decalcified with EDTA (Sigma-Aldrich, St. Louis, MO, USA) and embedded in paraffin (Sigma-Aldrich, St. Louis, MO, USA), and sections were stained with H&E (Sigma-Aldrich, St. Louis, MO, USA). Slides were scanned using Aperio AT Turbo (Aperio, Vista, CA, USA) and analysed using Aperio ImageScope v10 software (Leica Biosystems, Waverley, Australia) and the Positive Pixel Count v9 algorithm.

### 2.11. Statistics

The *t*-test was used when the differences in variances was <4 fold. Otherwise, the non-parametric Kolmogorov-Smirnov exact test was used (GraphPad Prism 10).

## 3. Results

### 3.1. Generation and characterisation of YYV-CHIKV_Mauritius_

A YYV-CHIKV_Mauritius_ chimera was generated using the CPER method through replacement of the YYV structural proteins with CHIKV_Mauritius_ structural proteins (Fig 1A). Replication kinetics in C6/36 mosquito cells between YYV-CHIKV_Mauritius_ and CHIKV_Mauritius_ found comparable titres, with YYV-CHIKV_Mauritius_ and CHIKV_Mauritius_ reaching peak titres at 48 hpi of 10^9.13^ TCID_50_/mL and 10^9.69^ TCID_50_/mL, respectively (Fig 1B).

**Figure 1.**
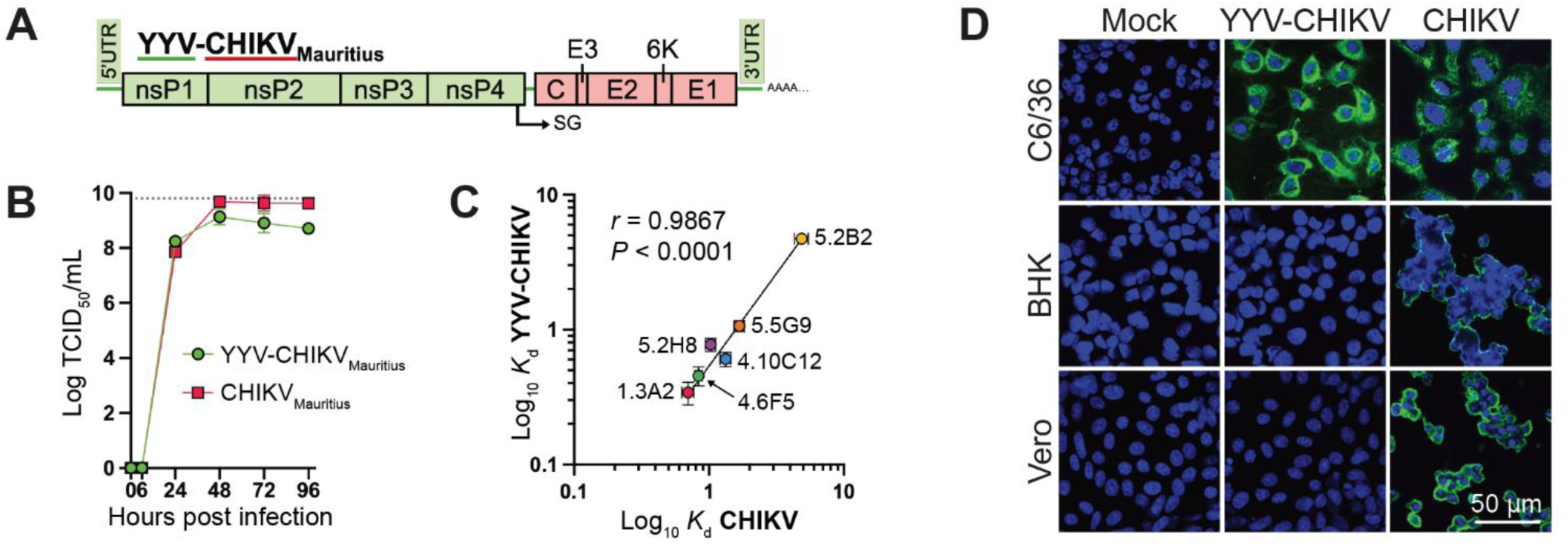
Generation and characterisation of YYV-CHIKV_Mauritius_. (A) Schematic of the YYV-CHIKV_Mauritius_ chimera, where the YYV genetic backbone (green) is displaying CHIKV_Mauritius_ structural protein genes (red). (B) Comparative growth kinetics between CHIKV_Mauritius_ and YYV-CHIKV_Mauritius_ in C6/36 cells, infected at an MOI of 0.1 (n = 3). Statistical analyses were performed with two-way ANOVA, and comparisons between individual time-points with Tukey’s multiple comparisons test. \*\**P* ≤ 0.01. (C) *K*_d_ values for six anti-CHIKV mAbs binding to YYV-CHIKV_Mauritius_ and CHIKV_Mauritius_ in fixed-cell ELISA. 1.3A2, 4.6F6, 4.10C12, 5.2B2, and 5.2H8 are E2 mAbs [ref, Lucas’ paper), and 5.5G9 is a capsid mAb [ref, Lucas’ other paper). Statistical analyses were performed with Pearson correlation. (D) IFA analysis of insect (C6/36) and mammalian (BSR and Vero) cell lines infected with YYV-CHIKV_Mauritius_ and CHIKV_Mauritius_ at an MOI of 1. Infected cell lines were probed with anti-CHIKV capsid mAb 5.5G9 and nuclei were stained with Hoechst 33342. Images taken at 40× magnification.

To investigate the antigenic authenticity of YYV-CHIKV_Mauritius_, a range of purified anti-CHIKV mAbs were tested in fixed-cell ELISA for YYV-CHIKV_Mauritius_ and CHIKV_Mauritius_. Comparing the apparent antibody dissociation constants (*K*_d_) found high correlation (r = 0.9933) between YYV-CHIKV_Mauritius_ and CHIKV_Mauritius_ binding kinetics, indicating strong antigenic authenticity of the chimera (Fig 1C).

We have previously demonstrated the inability for a YYV chimera with RRV structural proteins to replicate in a range of vertebrate cells (Bell et al., 2024). To confirm this same phenotype with YYV-CHIKV_Mauritius_, and to produce key data underpinning the safety of this virus, similar vertebrate-cell infection assays were required to be completed. C6/36, Vero and BSR cells were infected with YYV-CHIKV_Mauritius_ and CHIKV_Mauritius_ at a MOI of 1, or mock-infected (Fig 1D). IFA of fixed coverslips at 5 dpi found YYV-CHIKV_Mauritius_ did not replicate in either vertebrate cell line, demonstrated by the lack of viral staining with a virus-specific mAb, despite clear replication in the C6/36 cell line. As expected, CHIKV_Mauritius_ readily infected both vertebrate cell lines and exhibited high levels of CPE.

Deep sequencing of purified YYV-CHIKV_Mauritius_ found no mutations in the CHIKV structural genes, however, four amino acid changes were found in the YYV non-structural proteins (Table S1). To investigate this further, the regions of suspected amino acid changes were sequenced via RT-PCR in the YYV infectious clone (Bell et al., 2024) (P_1_) and the YYV-CHIKV_Mauritius_ stock (P_3_) used to the prepare the purified prep. These results found all mutations present in the parental YYV-CHIKV_Mauritius_ stock and one present in the YYV infectious clone (Ala387Val). Ala387Val was also present in raw sequencing data available on GenBank (SRR12113258), indicating this is from a YYV quasi-species. The other three mutations are likely compensatory or adaptive mutations resulting from continuous passaging in C6/36 cells, and do not permit YYV-CHIKV_Mauritius_ to infect vertebrate cells (Fig 1D).

To investigate whether YYV chimeric viruses may be used in surrogate neutralisation assays in the same way as flavivirus chimeric viruses (Hobson-Peters et al., 2019), the performance of YYV-CHIKV_Mauritius_ was compared to CHIKV_Mauritius_ in the microneutralisation format. Three neutralising anti-CHIKV E2 mAbs were assessed against YYV-CHIKV_Mauritius_ and CHIKV_Mauritius_ in C6/36 cells. These mAbs neutralised wild-type CHIKV and chimeric CHIKV to similar end-point titres (within 2-fold, Table 1). These results indicate that YYV-CHIKV_Mauritius_ may be used as a diagnostic antigen to detect neutralising CHIKV antibodies in human or animal sera, which will be investigated in future studies.

**Table 1:** Activity of neutralising E2 mAbs between chimera and wild type virus.

|  |  |  |
| --- | --- | --- |
| Antibody | Amount of antibody (µg) required for 100% virus neutralisation<sup>a</sup> |  |
|  | YYV-CHIKV<sub>Mauritius</sub> | CHIKV<sub>Mauritius</sub> |
| 1.3A2 | 10 | 5 |
| 4.6F5 | 2.5 | 2.5 |
| 4.10C12 | 10 | 10 |
| 5.5G9<sup>b</sup> | >10 | >10 |
| 4G2<sup>c</sup> | >10 | >10 |
| <sup>a</sup> Determined microscopically via CPE presence in two replicate wells per antibody dilution<br><sup>b</sup> 5.5G9 is a non-neutralising CHIKV capsid mAb, used as control mAb<br><sup>c</sup> 4G2 is a neutralising cross-reactive flavivirus envelope mAb, used as control mAb |  |  |

### 3.2. YYV-CHIKV_Mauritius_ provided immunogenicity in mice, and complete protection against CHIKV challenge

To determine whether YYV-CHIKV_Mauritius_ could induce a protective immune response against CHIKV_Reunion_ infection, groups of adult 6 week old female C57BL/6J mice were immunised with 2 doses of 1 µg of YYV-CHIKV_Mauritius_ or PBS and antibody responses and protection against disease and challenge were evaluated (Fig 2A). After one vaccination of 1 µg of YYV-CHIKV_Mauritius_, CHIKV-specific antibodies were observed by enzyme-linked immunosorbent assay (ELISA), which were significantly increased after a booster immunisation (Fig 2B). No CHIKV-specific antibodies were observed by ELISA for PBS vaccinated mice. Vaccination with 1 µg of YYV-CHIKV_Mauritius_ did not induce detectable CHIKV neutralising antibodies, however, a booster immunisation provided a significant increase neutralising antibody responses, with sera from 11 out of 12 mice able to neutralising CHIKV_Reunion_ (Fig 2C).

**Figure 2.**
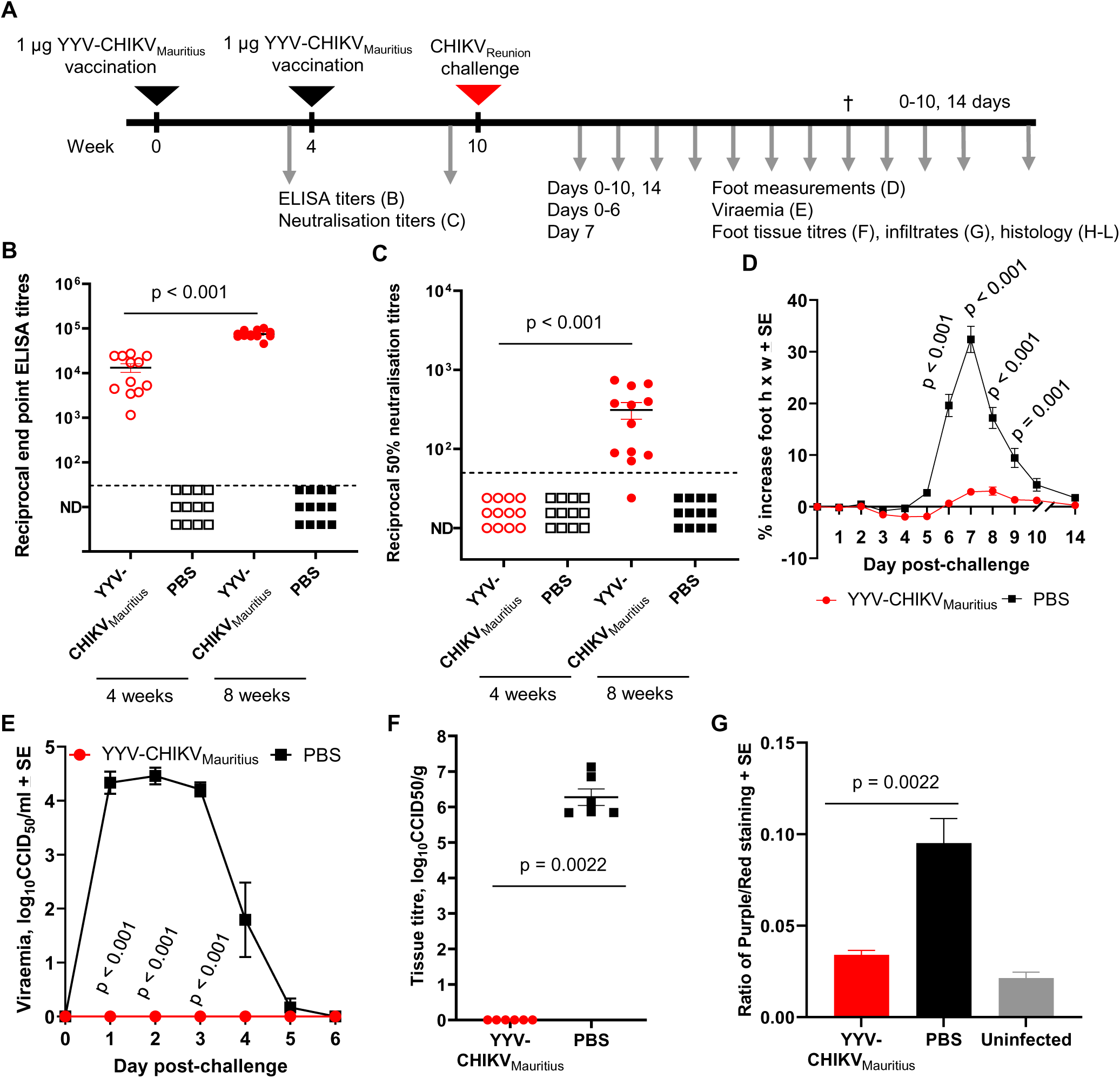
Evaluation of YYV-CHIKV_Mauritius_ in adult C57BL/6J mice after challenge with CHIKV Reunion. (A) Timeline of vaccination with YYV-CHIKV_Mauritius_ or PBS, with antibody measurements after single or double immunisations, followed by CHIKV_Reunion_ challenge, and subsequent disease determinations. (B) CHIKV_Reunion_ endpoint IgG ELISA titres after 1 or 2 vaccinations of female 6 week old C57BL/6J adult mice with non-adjuvated YYV-CHIK_Mauritius_ or PBS control. Dotted lines indicate a limit of detection (1:30 serum dilution). Statistical analysis used the Kolmogorov-Smirnov test. Lines among the data points indicate average and standard errors (C) CHIKV_Reunion_ 50% neutralising titres after 1 or 2 vaccinations of female 6 week old C57BL/6J adult mice with non-adjuvated YYV-CHIKV_Mauritius_ or PBS control. Dotted lines indicate a limit of detection (1:50 serum dilution). Statistical analysis used the Kolmogorov-Smirnov test. Lines among the data points indicate average and standard errors. (D) Percentage increase in foot height × width (relative to day 0) for C57BL/6J mice vaccinated as described for panel B, with *n* = 24 feet from 12 mice per group per time point from days 0-7; with n=12 feet from 6 mice per group per time point from days 8 onwards. Statistical analysis was with the *t* test. (E) CHIKV_Reunion_ viremia post challenge in mice vaccinated twice with non-adjuvanted YYV-CHIK_Mauritius_ or PBS control (*n* = 12 mice per group). The limit of detection for each mouse was 10^2^ TCID_50_/mL, with means from 12 mice plotted. Statistical analysis was with the Kolmogorov-Smirnov test. (F) CHIKV_Reunion_ tissue titres at day 7 post challenge in mice vaccinated twice with non-adjuvanted YYV-CHIK_Mauritius_ or PBS control (*n* = 6 mice per group). The limit of detection for each mouse was 10^2^ TCID_50_/mL, with means from 6 mice plotted. (G) Ratio of nuclear (purple) to nonnuclear (red) staining of H&E-stained foot sections (*n* = 6 mice, 6 feet per group, 3 sections per foot; values were averaged to produce one value for each foot). Statistical analysis used the Kolmogorov-Smirnov test.

To evaluate protection against CHIKV disease, mice were challenged with CHIKV_Reunion_ six weeks after the second YYV-CHIKV_Mauritius_ vaccination (Fig 2A). A challenge dose of 2 x 10^4^ CCID_50_ was used, as per our established model (Gardner et al., 2010; Metz et al., 2013; Nguyen et al., 2020; Poo et al., 2014) which shows overt foot swelling, characteristic cellular infiltrates and disease features that recapitulated CHIKV human disease. Two doses of 1 µg of YYV-CHIKV_Mauritius_ provided significant protection against foot swelling compared to the PBS group from days 6-9 (Fig 2D). Furthermore, mice vaccinated with two doses of YYV-CHIKV_Mauritius_ showed complete protection against viraemia (Fig 2E) and viral feet tissue titres (Fig 2F) compared to the PBS control group. Thus, two doses of 1 µg non-adjuvanted YYV-CHIKV_Mauritius_ generated ELISA responses and neutralising antibodies sufficient for complete protection against CHIKV infection.

### 3.3. YYV-CHIKV_Mauritius_ provided complete protection against CHIKV histopathology

To evaluate protection against CHIKV histopathology, H&E staining of mice feet at day 7 (time point of peak foot swelling), was performed. Quantification using purple (nuclear) versus red (cytoplasmic) staining ratios (Fig 2G), demonstrated significant differences between the YYV-CHIKV_Mauritius_ vaccinated mice compared to the PBS vaccinated mice. This indicated that there were significantly less cellular infiltrates in YYV-CHIKV_Mauritius_ vaccinated mice compared to the control vaccinated mice. No significant differences were observed between the YYV-CHIKV_Mauritius_ vaccinated mice and the uninfected controls.

H&E staining of feet from mice at 7 days post-challenge (Fig 3A) illustrated the characteristic mononuclear cellular infiltrates evident in muscle tissues (Fig 3B), in tendons (Fig 3C), in surrounding joint areas (Fig 3D) and in regions of subcutaneous oedema in the PBS-vaccinated control group (Fig 3E, third column). Haemorrhage was also observed in the PBS-vaccinated control group (Fig 3F, third column). Mice vaccinated with YYV-CHIKV_Mauritius_ demonstrated no evidence of cellular infiltrates in these regions (Fig 3, second column), with no major differences observed with healthy uninfected controls (Fig 3, first column). Thus, two doses of YYV-CHIKV_Mauritius_ provided complete protection against CHIKV myositis, tendonitis, arthritis and haemorrhage.

**Figure 3.**
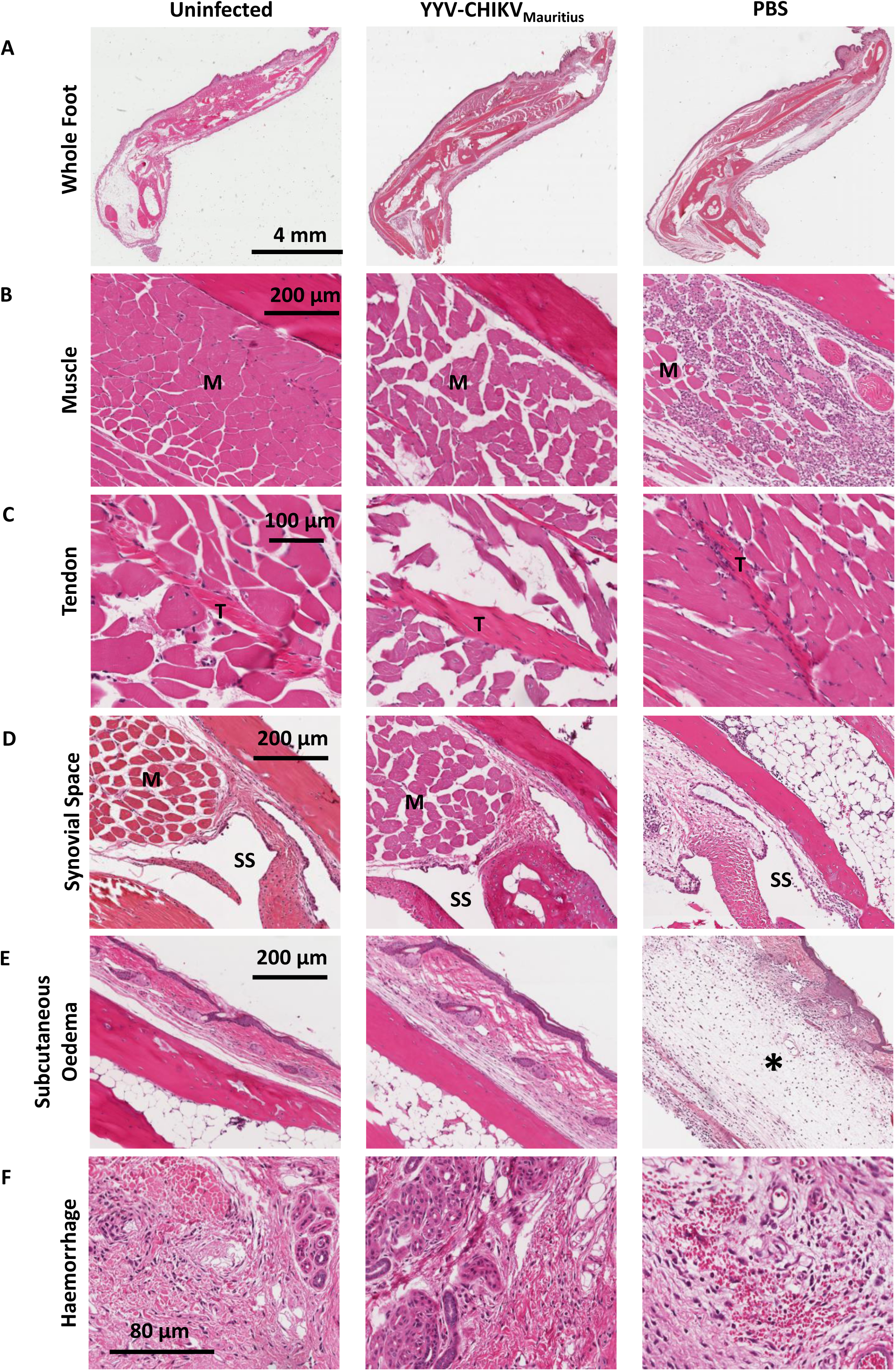
Histopathology of YYV-CHIKV_Mauritius_ vaccinated adult C57BL/6J mice after challenge with CHIKV Reunion. (A) H&E staining of whole feet of mouse feet 7 days post-challenge. (B) H&E staining of muscle tissues in foot sections of mice vaccinated twice with nonadjuvanted YYV-CHIK_Mauritius_ or PBS control. M = muscle. (C) As for (B) with tendons. (D) As for (B) with joint areas. (E) as for (B) with subcutaneous oedema regions. (F) As for (B) with haemorrhagic regions.

### 3.4. YYV-RRV_TT_ provided immunogenicity in mice, and complete protection against RRV challenge, while YYV-CHIKV_Mauritius_ provided some immunogenicity in mice but no cross-protection against RRV challenge

To determine whether YYV-RRV_TT_ could induce a protective immune response against RRV_TT_ infection, groups of adult 6 week old female C57BL/6J mice were immunised with 2 doses of 1 µg of YYV-RRV_TT_ or PBS and antibody responses and protection against disease and challenge were evaluated (Fig 4A). Mice were also vaccinated with YYV-CHIKV_Mauritius_ to determine potential cross-protection against RRV_TT_ challenge (Fig 4A).

**Figure 4.**
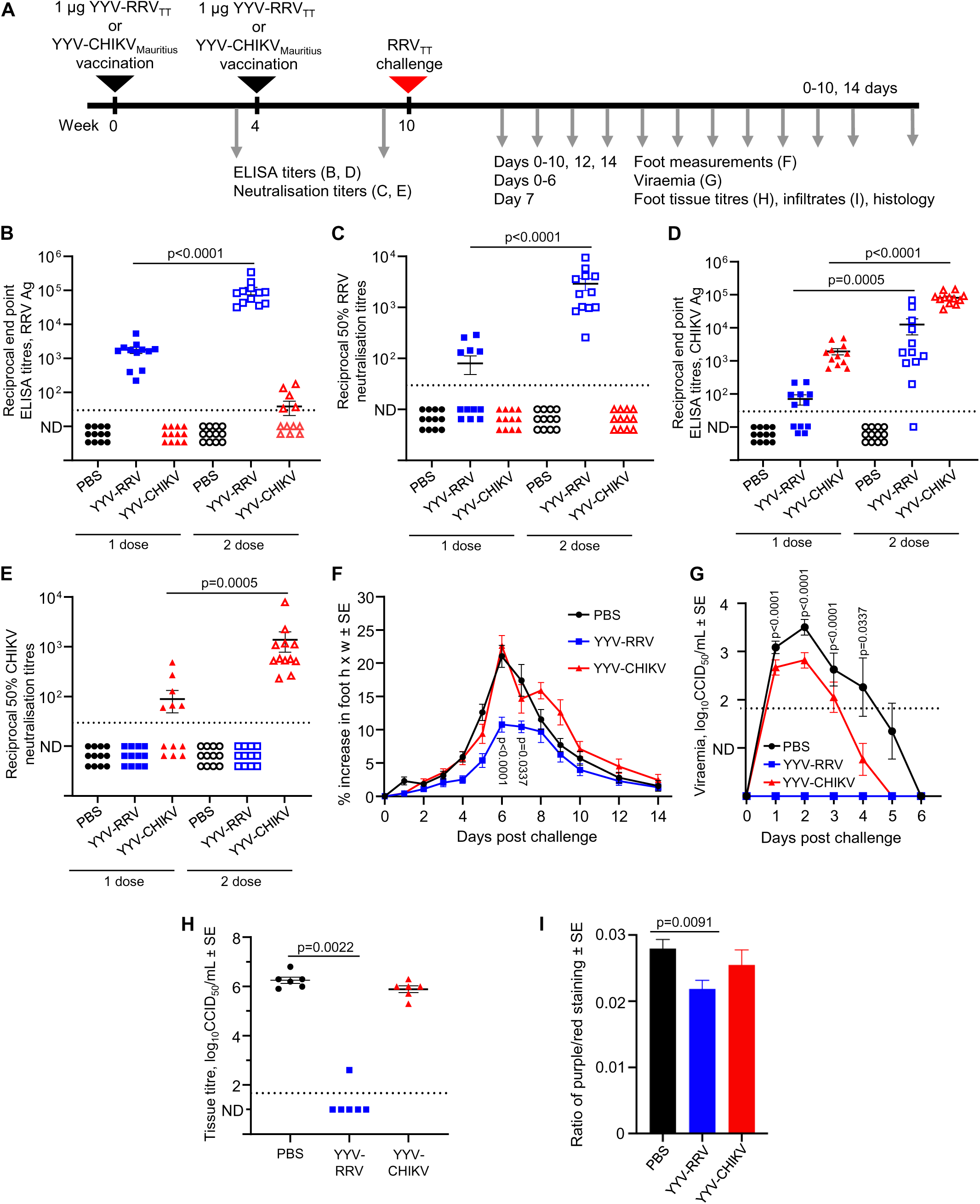
Evaluation of YYV-RRV_TT_ protection and YYV-CHIKV_Mauritius_ cross-protection against RRV_TT_ challenge in adult C57BL/6J mice. (A) Timeline of vaccination with YYV-RRV_TT_, YYV-CHIKV_Mauritius_, or PBS, with antibody measurements after single or double immunisations, followed by RRV_TT_ challenge, and subsequent disease determinations. (B) RRV_TT_ endpoint IgG ELISA titres after 1 or 2 vaccinations of female 6 week old C57BL/6J adult mice with non-adjuvanted YYV-RRV_TT_, YYV-CHIKV_Mauritius_, or PBS. Dotted lines indicate a limit of detection (1:30 serum dilution). Statistical analyses by Kolmogorov-Smirnov test. Lines among the data points indicate average and standard errors. (C) RRV_TT_ 50% neutralising titres after 1 or 2 vaccinations of female 6 week old C57BL/6J adult mice with non-adjuvanted YYV-RRV_TT_, YYV-CHIKV_Mauritius_, or PBS. Dotted lines indicate a limit of detection (1:40 serum dilution). Statistical analysis used the Kolmogorov-Smirnov test. Lines among the data points indicate average and standard errors. (D) CHIKV_Reunion_ endpoint IgG ELISA titres after 1 or 2 vaccinations of mice as described in (B). Dotted lines indicate a limit of detection (1:30 serum dilution). Statistical analysis by Kolmogorov-Smirnov test. Lines among the data points indicate average and standard errors (E) CHIKV_Reunion_ 50% neutralising titres after 1 or 2 vaccinations of female 6 week old C57BL/6J adult mice as described in (C). Dotted lines indicate a limit of detection (1:40 serum dilution). Statistical analysis by Kolmogorov-Smirnov test. Lines among the data points indicate average and standard errors. (F) Percentage increase in foot height × width (relative to day 0) for C57BL/6J mice vaccinated as described in (A), with *n* = 24 feet from 12 mice per group per time point from days 0-6; with n=12 feet from 6 mice per group per time point from days 7 onwards. Statistical analysis by *t* test. (G) RRV_TT_ viremia post challenge in mice as described in (A) (*n* = 12 mice per group). The limit of detection for each mouse was 10^2^ CCID_50_/ml, with means from 12 mice plotted. Statistical analysis by Kolmogorov-Smirnov test. (H) RRV_TT_ tissue titres at day 6 post challenge in mice vaccinated twice with non-adjuvanted as described in (A) (*n* = 6 mice per group). The limit of detection for each mouse was 10^2^ CCID_50_/mL, with means from 6 mice plotted. (I) Ratio of nuclear (purple) to nonnuclear (red) staining of H&E-stained foot sections (*n* = 6 mice, 6 feet per group, 3 sections per foot; values were averaged to produce one value for each foot). Statistical analysis by t-test Welch’s correction).

After one vaccination of 1 µg of YYV-RRV_TT_, RRV-specific antibodies were observed by ELISA, which were significantly increased after a booster immunisation (Fig 4B). No RRV-specific antibodies were observed by ELISA for PBS or YYV-CHIKV_Mauritius_ vaccinated mice after one vaccination. However, cross-reactive RRV-specific antibodies were detected in 5 out of 12 mice after a booster immunisation with YYV-CHIKV_Mauritius_. Vaccination with 1 µg of YYV-RRV_TT_ induced RRV neutralising antibodies in 5 out of 12 mice with a significant increase in neutralising antibodies after a second vaccination (Fig 4C). Vaccination with PBS or YYV-CHIKV_Mauritius_ did not induce any RRV neutralising antibodies after one dose, or a booster dose (Fig 4C). Reciprocally, one vaccination of 1 µg of YYV-RRV_TT_ induced cross-reactive CHIKV-specific antibodies in 7 out of 12 mice, which was significantly increased after a second dose (Fig 4D). As expected, CHIKV-specific antibodies were observed by ELISA in mice vaccinated with one dose of YYV-CHIKV_Mauritius_ which was significantly increased after a booster vaccination (Fig 4D). No CHIKV neutralising antibodies were detected in PBS or YYV-RRV_TT_ vaccinated mice after one or two immunisations (Fig 4E). CHIKV neutralising antibodies were detected in 50% of YYV-CHIKV_Mauritius_ vaccinated mice, which was significantly increased after a second immunisation with all mice having CHIKV neutralising antibodies (Fig 4E).

To evaluate protection against RRV disease, mice were challenged with RRV_TT_ six weeks after the second YYV-CHIKV_Mauritius_ vaccination (Fig 4A). A challenge dose of 2 x 10^4^ CCID_50_ was used, as per our established model (Nguyen et al., 2020) which shows foot swelling, characteristic cellular infiltrates and disease features that recapitulated RRV human disease. Two doses of 1 µg of YYV-RRV_TT_ provided significant protection against foot swelling compared to the PBS and YYV-CHIKV_Mauritius_ group from days 6-7 (Fig 4F). Furthermore, mice vaccinated with two doses of YYV-RRV_TT_ showed complete protection against viraemia (Fig 4G), and protection against viral feet tissue titres with 5 out of 6 mice having complete protection (Fig 4H), significantly different compared to the PBS and YYV-CHIKV_Mauritius_ vaccinated groups. Thus, two doses of 1 µg non-adjuvanted YYV-RRV_TT_ generated ELISA responses and neutralising antibodies sufficient for complete protection against RRV_TT_ viraemia and arthritic disease. In comparison, YYV-CHIKV vaccinated mice demonstrated some cross-immunogenicity, but offered no cross-protection against RRV_TT_.

### 3.5. YYV-RRV_TT_ provided complete protection against RRV histopathology

To evaluate protection against RRV histopathology, H&E staining of mice feet at day 6 (time point of peak foot swelling), was performed and analysed. Quantification using purple (nuclear) versus red (cytoplasmic) staining ratios (Fig 4I), demonstrated significant differences between the YYV-RRV_TT_ vaccinated mice compared to the PBS vaccinated mice. This indicated that there were significantly less cellular infiltrates in YYV-CHIKV_Mauritius_ vaccinated mice compared to the PBS vaccinated mice. No significant differences were observed between the YYV-RRV_TT_ vaccinated and YYV-CHIKV_Mauritius_ vaccinated mice, or the PBS vaccinated and YYV-CHIKV_Mauritius_ vaccinated mice.

H&E staining of feet from mice at 6 days post-challenge (Fig 5A) illustrated the characteristic mononuclear cellular infiltrates evident in muscle tissues (Fig 5B), in tendons (Fig 5C), in surrounding joint areas (Fig 5D) and in regions of subcutaneous oedema in the PBS-vaccinated and YYV-CHIKV_Mauritius_ vaccinated group (Fig 5E, second column, and Fig 5E, last column, respectively). Mice vaccinated with YYV-RRV_TT_ demonstrated no evidence of cellular infiltrates in these regions (Fig 5E, third column), with no major differences observed with healthy uninfected controls (Fig 5E, first column). Thus, two doses of YYV-RRV_TT_ provided complete protection against RRV myositis, tendonitis and arthritis.

**Figure 5.**
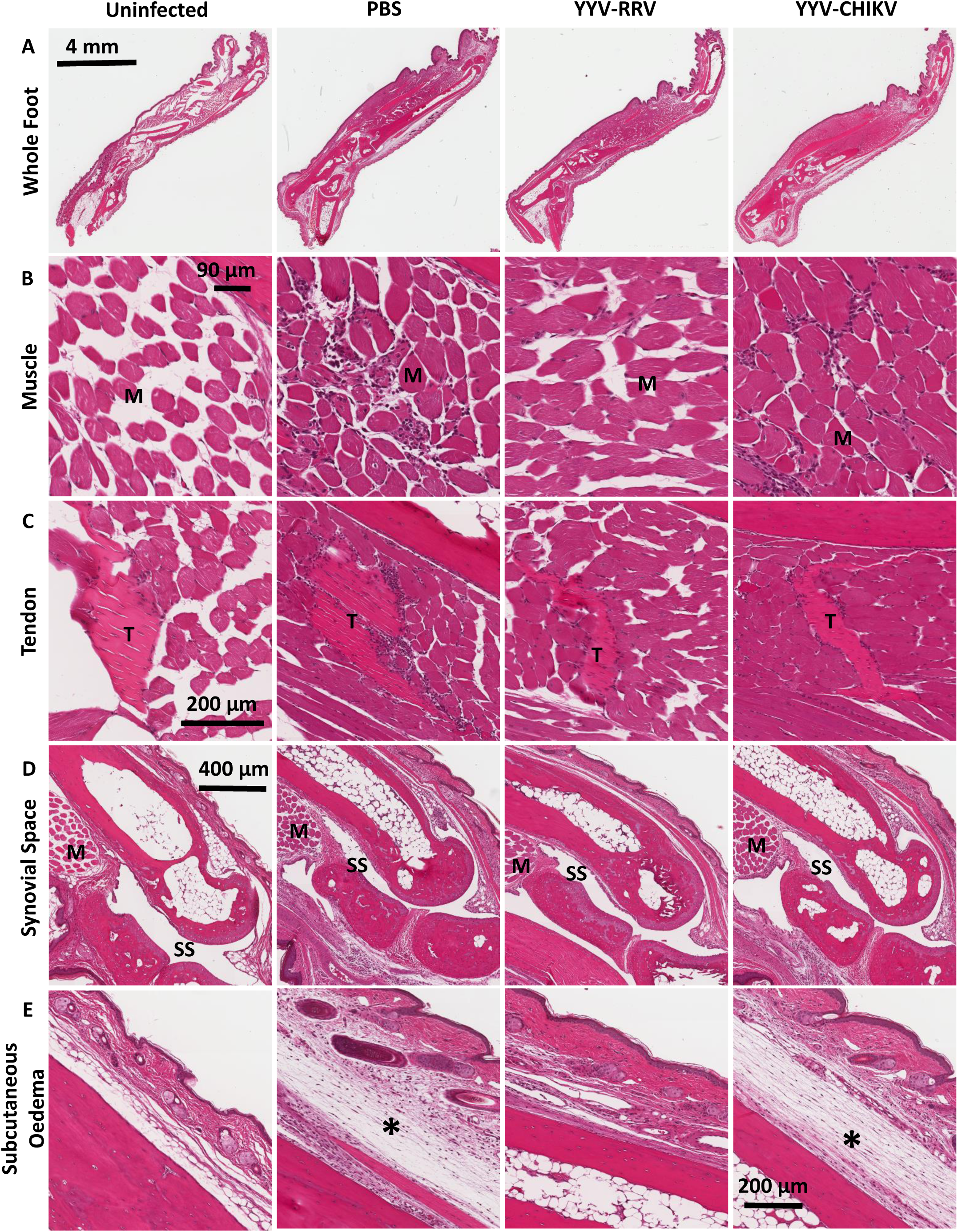
Histopathology of YYV-RRV_TT_ compared to YYV-CHIKV_Mauritius_ and PBS vaccinated adult C57BL/6J mice after RRV_TT_ challenge. (A) H&E staining of whole feet of mouse feet 6 days post-challenge. (B) H YYV-RRV_TT_, YYV-CHIKV_Mauritius_, or PBS. M = muscle. (C) As for (B) with tendons. (D) As for (B) with joint areas. (E) as for (B) with subcutaneous oedema regions.

## 4. Discussion

Since its initial isolation in 1952, CHIKV has caused widespread outbreaks in more than 100 countries over four continents, resulting in >10 million cases of debilitating rheumatic disease. The virus has continued to circulate in endemic regions of Asia and South America, with more than 75% of the world population estimated to be at risk of CHIKV infection (Bettis et al., 2022). Two CHIKV vaccines have recently received FDA approval. IXCHIQ® (Valneva), a live-attenuated vaccine, was approved in 2023 and showed strong immunogenicity, with nearly all participants developing neutralising antibodies within a month (U.S. Food & Drug Administration, 2023). VIMKUNYA™ (PXVX0317, Bavarian Nordic), a virus-like particle (VLP) vaccine approved in 2025 for individuals aged 12 and older, also demonstrated robust and sustained antibody responses in phase III trials (Bavarian Nordic, 2025). While highly immunogenic, live-attenuated vaccines pose increased safety concerns compared to most other vaccine technologies, and are not suitable for immunocompromised or pregnant individuals.

Insect-specific vaccines are alternative and safe approaches for the development of whole-virus vaccines as they possess multiple barriers restricting their replication in vertebrates (Nasar et al., 2015a; Nasar et al., 2015b). EILV, another insect-specific alphavirus, has previously been explored to generate an EILV/CHIKV-sP chimeric vaccine (Adam et al., 2021; Adam et al., 2024). These studies demonstrated that a single unadjuvanted dose of 10^8.1^ PFU of EILV/CHIKV-sP (live virus, PEG-precipitated) protected mice and non-human primates from CHIKV challenge, with a subsequent study demonstrating that a dose of 10^8^ PFU was the minimal optimal single dose to generate strong antibody responses in mice (Adam et al., 2024). This dose is more than the approved IXCHIQ® vaccine which only contains 10^4^ TCID_50_ of the live-attenuated virus in a single 0.5 mL dose (U.S. Food & Drug Administration, 2023).

Herein, we describe the application of the insect-specific alphavirus YYV platform technology for the development of the YYV-CHIKV_Mauritius_ chimeric vaccine as a safer alternative CHIKV vaccine candidate and demonstrate its efficacy in protecting against CHIKV arthropathies in a wild-type mouse model of CHIKV infection and disease. A single unadjuvanted dose of 1 µg was sufficient to generate anti-CHIKV ELISA responses, although a second dose was required for neutralising antibody responses to be detected in 11/12 mice. This regime was sufficient to protect vaccinated mice against foot swelling, viraemia, feet tissue titres and histological features of CHIKV disease including myositis, tendonitis, subcutaneous oedema, haemorrhage and arthritis. Both EILV and YYV vaccine platforms show promise in generating chimeric vaccines against CHIKV (and several other arbovirus pathogens), but a direct comparison of efficacy cannot be easily achieved due to differences in composition and manufacturing.

We also describe the application and development of the YYV-RRV_TT_ chimeric vaccine as a safe vaccine candidate which demonstrates efficacy in protecting against RRV arthropathies in a wild-type RRV mouse model of infection and disease. A single unadjuvanted dose of 1 µg was sufficient to generate anti-RRV ELISA responses, with 5/12 mice already demonstrating neutralising antibodies. A booster immunisation showed significantly higher anti-RRV ELISA responses with all mice having over neutralising antibodies. This regime was sufficient to protect vaccinated mice against foot swelling, viraemia, viral feet tissue titres and histological features of RRV disease including myositis, tendonitis, subcutaneous oedema, and arthritis. With no additional progression from the UV and formalin-inactivated RRV vaccine currently owned by Resilience Government Services, Inc, YYV-RRV_TT_ may be a viable vaccine candidate for progression to clinical trials given the advantages of the insect-specific virus vaccine platform including scalability, cost and immunogenicity. In addition, with ∼7.63 million international visitors to Australia in 2024 (Australian Trade and Investment Commission, 2025), this may provide incentive to market and commercialise an RRV travel vaccine. The expected increasing burden of RRV infection attributable to increasing temperatures and climate change underscores the need to develop therapeutics (Damtew et al., 2024).

We also evaluated the potential for cross-protection against RRV using YYV-CHIKV_Mauritius_, as a CHIKV vaccine would be deemed more commercially viable internationally. Although suboptimal cross-reactive RRV ELISA antibodies were detected in some mice after two immunisations with YYV-CHIKV_Mauritius_, no neutralising antibodies were detected. Subsequently, no protection against RRV foot swelling, viraemia, viral feet tissue titres or histopathological lesions was observed. Although CHIKV and RRV belong to the same Semliki Forest complex serogroup, many studies have demonstrated that cross-protection, while possible, is often not complete nor universal for all virus combinations within the serogroup (Abbo et al., 2023; Gardner et al., 2010; Nguyen et al., 2020; Schmidt et al., 2022). For efficient cross-protection, much higher immune responses induced by an elevated vaccine dose, additional booster immunisations, or changes to the presenting antigen would be required.

IgG and IgM antibody detection is routinely used for serological alphavirus diagnosis. However, antibodies may cross-react to other alphaviruses, which can cause problems for specific diagnosis especially in regions where multiple alphaviruses potentially co-circulate, although the extent and effect of this behaviour is not fully understood. Herein, we show that YYV-CHIKV_Mauritius_ chimeras are antigenically identical to their wild-type counterparts, supported by published data on EILV chimeras (Adam et al., 2021; Adam et al., 2024). These data demonstrate the potential utility of YYV chimeras as safe diagnostic antigens, which may be used in lieu of pathogenic wild-type virus. Further studies will thoroughly investigate the use of YYV chimeras in diagnostics, including application to alternative methods such as MIAs and point-of-case lateral flow devices, which have successfully been used in complex with ISF chimeras (Hobson-Peters et al., 2019; Johnston et al., 2025). In conclusion, we show that YYV-CHIKV_Mauriitus_ is a promising insect-specific alphavirus chimeric vaccine platform which protects against CHIKV infection and disease in a mouse model of CHIKV arthropathy. We also show that YYV-RRV_TT_ is a viable vaccine candidate which protects against RRV infection and disease in a mouse model of RRV arthropathy.

## Acknowledgements

We would like to thank Madeline Thompson for techical assistance with the YYV-CHIKV prep. We thank the QIMR Berghofer animal facility, histology facility, microscopy facility, and PC3 Facility Manager.

