## Supplementary Table 1 for "Insect-specific Yada Yada virus chimeric vaccines protect against chikungunya and Ross River virus-induced arthritis"

**Table S1:** Deep sequencing of YYV-CHIKV$_{Mauritius}$.

| Gene | Mutation | Nucleotide | Present in mosquito pool sequencing |
|---|---|---|---|
| nsP1 | Leu52Phe | TTG → TT**T** | No |
| nsP3 | Ala75Ser | GCT → **T**CT | No |
| nsP3 | Arg327Ser | CGT → **A**GT | No |
| nsP3 | Ala387Val | GCA → G**T**A | Yes (SRR12113258) |
